# A bioorthogonal chemical reporter for fatty acid synthase-dependent protein acylation

**DOI:** 10.1101/2021.05.07.443132

**Authors:** Krithika P. Karthigeyan, Lizhi Zhang, David R. Loiselle, Timothy A. J. Haystead, Menakshi Bhat, Jacob S. Yount, Jesse J. Kwiek

## Abstract

Cells acquire fatty acids from dietary sources or via *de novo* palmitate production by fatty acid synthase (FASN). Although most cells express FASN at low levels, it is upregulated in cancers and during replication of many viruses. The precise role of FASN in disease pathogenesis is poorly understood, and whether *de novo* fatty acid synthesis contributes to host or viral protein acylation has been traditionally difficult to study. We describe a cell permeable, click-chemistry compatible alkynyl-acetate analog (Alk-4) that functions as a reporter of FASN-dependent protein acylation. In a FASN-dependent manner, Alk-4 selectively labeled the cellular protein interferon-induced transmembrane protein 3 (IFITM3) at its palmitoylation sites, and the HIV-1 matrix protein at its myristoylation site. Alk-4 metabolic labeling also enabled biotin-based purification and identification of more than 200 FASN-dependent acylated cellular proteins. Thus, Alk-4 is a useful bioorthogonal tool to selectively probe FASN-mediated protein acylation in normal and diseased states.

## Introduction

Long chain fatty acids (FA) are essential components of lipid bilayers, are used to store energy liberated by β-oxidation, and are covalently attached to proteins to increase hydrophobicity and regulate subcellular localization.^1^ Long chain fatty acids can be obtained exogenously through dietary sources, or endogenously via *de novo* fatty acid biosynthesis.^2^ Mammalian fatty acid synthase (FASN) is a 272 kDa cytosolic enzyme that catalyzes the complete *de novo* synthesis of palmitate from acetyl-CoA and malonyl-CoA. The final product, palmitic acid (16:0) is then released from FASN, where it can be metabolized by β-oxidation into myristic acid (14:0), or other long chain FA.^3^ FASN expression is highly regulated in cells and its expression can change dramatically in response to stresses such as starvation, lactation or pathological states.^3^ Increased *de novo* FA biosynthesis and FASN up-regulation have been observed in breast cancer, melanoma, and hepatocellular carcinoma.^4^ Studies of enveloped viruses including hepatitis B virus,^5^ Dengue virus,^6^ Epstein-Barr virus,^7^ hepatitis C virus,^8^ HIV-1,^9^ Chikungunya virus,^10,11^ and West Nile virus^12,13^ indicate that many viruses both upregulate and require host FASN activity for effective replication. The contributions of *de novo* synthesized FA to post-translational modifications of viral and host proteins remains understudied.

Identification of protein acylation has been challenging due to the lack of antibodies against lipid modifications, and inefficiencies of standard mass spectrometry techniques to identify acylated proteins.^14^ While protein myristoylation site prediction is facilitated by a consensus sequence motif on nearly all myristoylated proteins (Met-Gly-XXX-Ser/Thr),^1^ protein palmitoylation site prediction remains challenging due to the lack of a consensus sequence.^15^ To measure acyl-group synthesis mediated by FASN and the fate of the *de novo* synthesized fatty acids, one must use ^14^C labeled acetate, which suffers from low detection sensitivity, general complications associated with radioisotope work,^16^ and an inability to selectively enrich acylated proteins. Over the last decade, bioorthogonal labeling and detection of protein fatty acylation using click chemistry compatible analogs of palmitate and myristate have provided quick and sensitive methods for detection of protein acylations.^17,18^ The copper-catalyzed azide-alkyne cycloaddition (CuAAC) reactions enable labeling of cells with alkynyl analogs of fatty acids that can be reacted with azides conjugated to suitable detection tags, such as fluorophores, or affinity tags, including biotin.^19,20^ Although very useful, palmitate and myristate analogs only measure the acylation state of proteins modified by the exogenous chemical reporters. Given the critical role of FASN-dependent *de novo* synthesized fatty acids in cancer, metabolic disorders, and viral replication, we posit that a bioorthogonal reporter of FASN-dependent protein acylation will facilitate a better understanding of the contributions of FASN-dependent protein fatty acylation to protein function, protein localization, and FASN-mediated pathogenesis. Here we demonstrate the utility of 5-hexynoic acid, or Alk-4, a cell permeable, click-chemistry compound that labels proteins acylated by products of FASN-mediated *de novo* fatty acid biosynthesis.

## Results

### Alk-4 labels palmitoylated proteins at known palmitoylation sites

Bioorthogonal reporters such as alkynyl palmitate (“Alk-16”) and alkynyl myristate (“Alk-12”) are substrates of palmitoyltransferase and myristoyltransferase activity that are often used to identify palmitoylated and myristoylated proteins.^21–23^ Because these reporters mimic the end product of FASN activity (palmitate) or palmitate oxidation (myristate), Alk-12 and Alk-16 cannot be used to determine the source of the fatty acyl adduct (i.e. exogenous/imported or endogenous/*de novo* synthesized). We hypothesized that a cell permeable, bioorthogonal mimic of a putative FASN substrate, 5-hexynoate (termed Alk-4 here)^24^ could be used to study the contributions of FASN-mediated *de novo* fatty acid synthesis to protein acylation (Figure 1a,b). To determine whether Alk-4 selectively labels palmitoylated proteins, we tested whether a known palmitoylated protein, IFITM3, was labeled upon a 24 hour treatment of cells with alkynyl acetate analogues of different carbon chain lengths, Alk-3 and Alk-4, in comparison with the well-established palmitoylation reporter Alk-16 (Figure 1a, b). Alk-16 robustly labeled IFITM3 as detected by click-chemistry tagging of immunoprecipitated IFITM3 with azido-rhodamine and fluorescence gel scanning. Alk-4 also successfully labeled IFITM3, while Alk-3 showed minimal labeling (Figure 2a). Next, we tested whether Alk-4 labeling of IFITM3 occurred on its known palmitoylated cysteines.^22^ A triple cysteine to alanine palmitoylation-deficient mutant of IFITM3 (termed PalmΔ), was not labeled by either Alk-16 or Alk-4 (Figure 2b). To further test the ability of Alk-4 to label palmitoylated proteins, we examined whether the tetraspanin CD9, which has 6 palmitoylated cysteines, was also labeled by Alk-4. Similar to IFITM3, CD9 was labeled by Alk-4, while a mutant CD9 in which its palmitoylated cysteines were mutated to alanine (termed CD9-PalmΔ), was not labeled (Figure 2c). These results indicate that Alk-4 is metabolized into a click-chemistry functionalized fatty acid adduct that is specifically incorporated onto protein palmitoylation sites.

**Figure 1:**
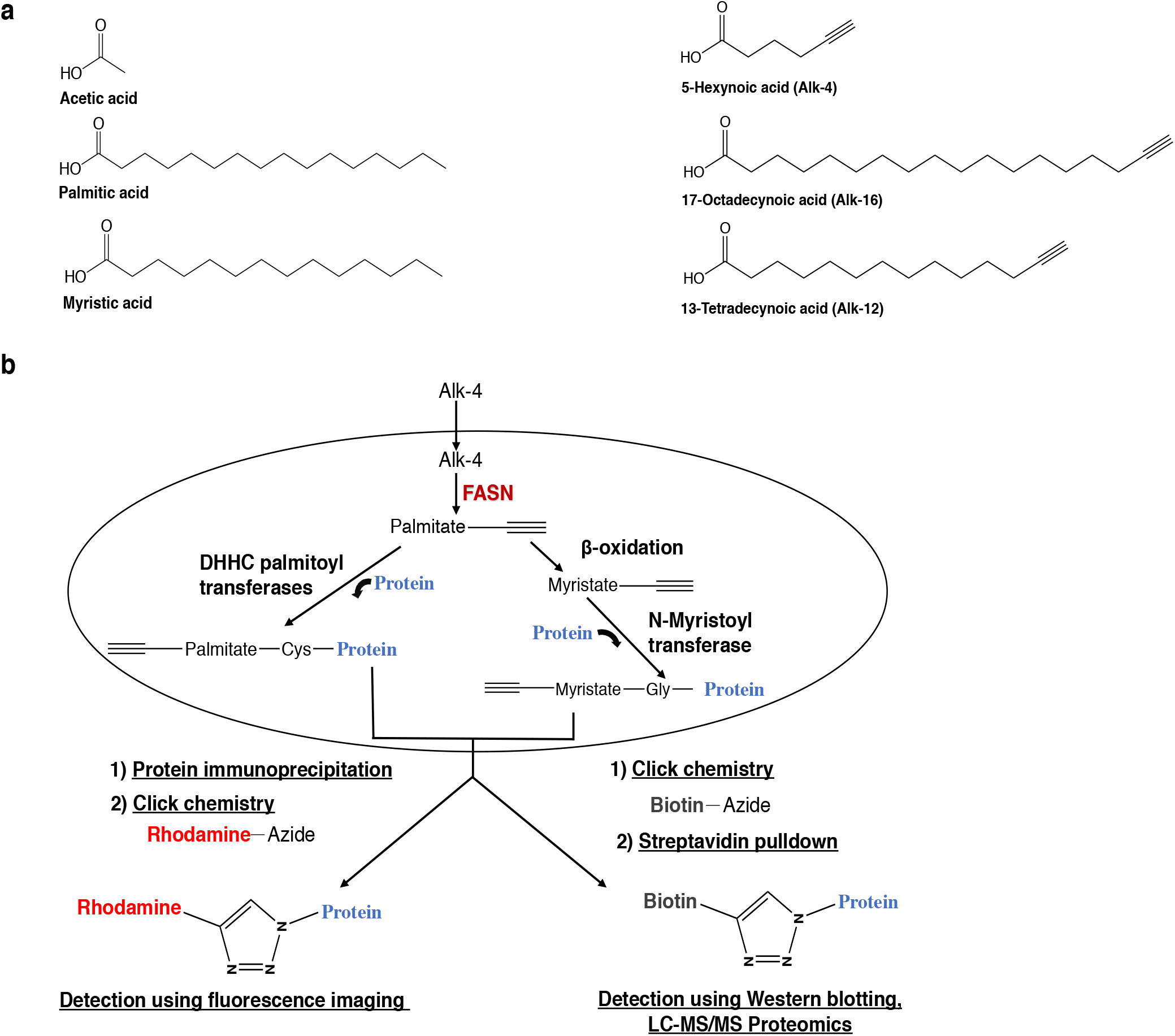
Analog structures and schema depicting Alk-4 metabolism and incorporation onto protein acylation sites. **(a)** Structures of acetate, palmitate, and myristate, followed by their click chemistry-compatible analogs Alk-4, Alk-16, and Alk-12. (**b)** 5-Hexynoic acid (Alk-4) can be metabolized through the endogenous FASN pathway to yield functionalized versions of fatty acyl groups that are transferred onto protein acylation sites by palmitoyl or N-myristoyl transferases. Copper-catalyzed azide-alkyne cycloaddition (CuAAC) click reaction of proteins containing the functionalized alkyne group to an azide-conjugated fluorophore such as rhodamine can be used for fluorescence imaging, while CuAAC click reaction of alkyne-containing proteins to an azide-conjugated biotin can be used for affinity purification and subsequent western blotting or proteomics.

**Figure 2:**
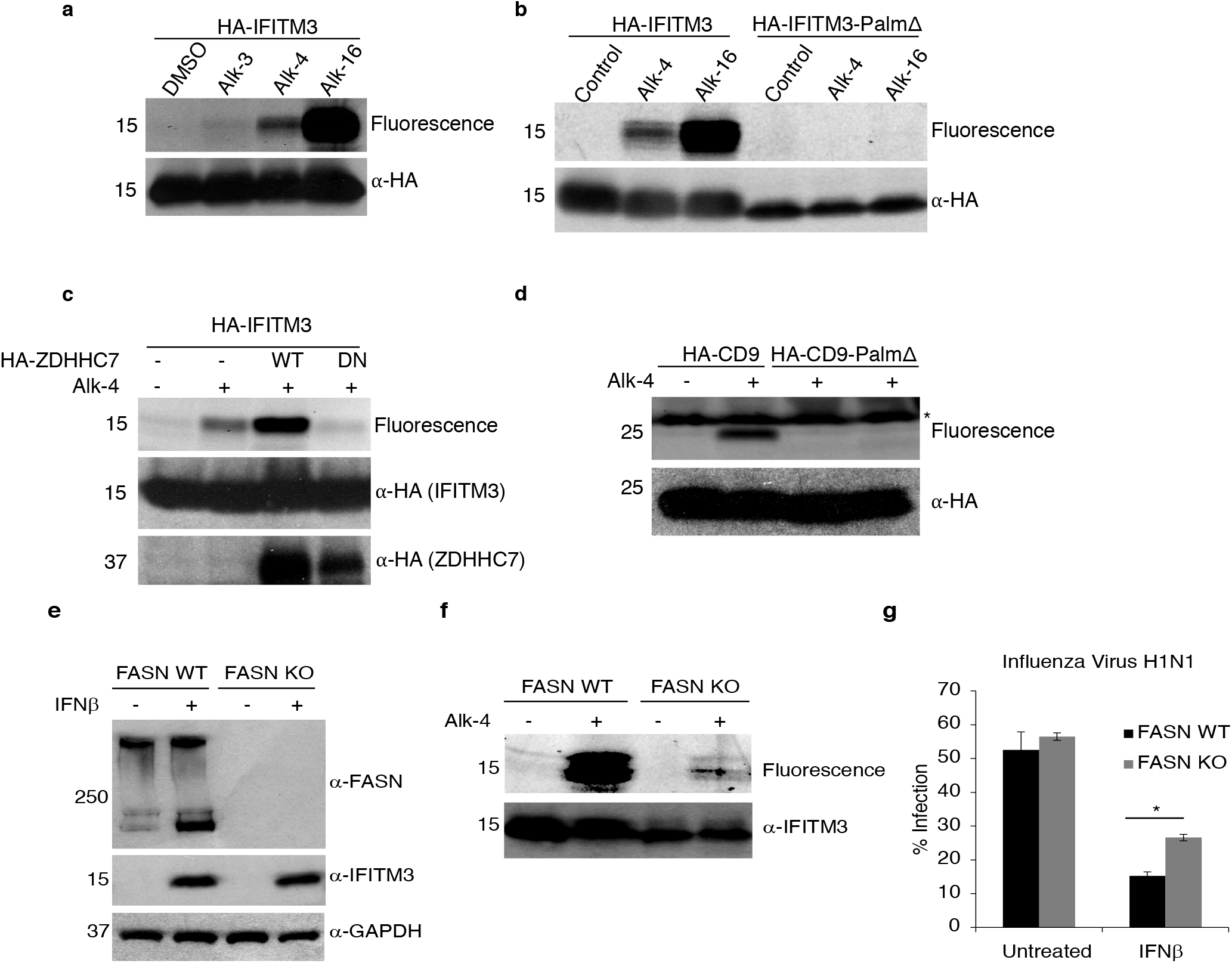
FASN-dependent incorporation of Alk-4 at known protein palmitoylation sites. **(a)** Immunoprecipitated HA-IFITM3 from HA-IFITM3 transfected 293Ts followed by rhodamine azide click reaction revealed detectable labeling by Alk-16 and Alk-4 and minimal labeling by Alk-3. **(b)** A triple cysteine to alanine IFITM3 mutant (Palm) was not labeled by Alk-4, suggesting that Alk-4 labeling of IFITM3 occurred on known palmitoylated cysteines. **(c)** Alk-4 labeled CD9, while a mutant where its six palmitoylated cysteines were mutated to alanine was not labeled, revealing that Alk-4 labels CD9 on known palmitoylated cysteines. **(d)** DHHC palmitoyltransferase overexpression increased Alk-4 labeling of IFITM3, while a dominant negative mutant partially decreased labeling, suggesting that Alk-4 is metabolized into a long chain fatty acid utilized by DHHC palmitoyltransferases (* indicates 25kDa fluorescent molecular weight standard bleed through). **(e)** To test the requirement of FASN for labeling of endogenous IFITM3 in cells treated with IFNβ, HAP1 WT and FASN KO cells were labeled with Alk-4. Western blotting to confirm FASN levels in WT and KO cells, and expression of endogenous IFITM3 on IFNβ treatment. **(f)** Alk-4 labeling of endogenous IFITM3 was only observed in WT cells and not detected in FASN KO cells, indicating that FASN contributes to palmitoylation of IFITM3. **(g)** IFNβ was significantly less effective at inhibiting Influenza virus strain H1N1 infection in FASN KO cells (*p = 0.0002), indicating that FASN is required for mounting of an effective IFNβ immune response against influenza virus, possibly through provision for fatty acyl groups for activation of IFITM3.

### Alk-4 metabolism provides a substrate used by DHHC palmitoyltransferases

Many proteins are reversibly palmitoylated at cysteine residues^25^ by aspartate-histidine-histidine-cysteine (DHHC) palmitoyltransferases. DHHC palmitoyltransferases primarily use palmitoyl-CoA (C16:0) to modify cysteine residues on proteins, although DHHC’s can tolerate substrates with carbon chain lengths as short as 14 and as long as 20.^26,27^ Acyl chains with fewer than 14 carbons have not been detected on cysteines, indicating that DHHC enzymes disfavor short chain fatty acids as substrates.^28–31^ Given the selectivity of the DHHC palmitoyltransferases for long chain fatty acids, we sought to determine whether labeling of IFITM3 by Alk-4 was affected by DHHC7 overexpression, which was previously shown to be among the enzymes that can catalyze IFITM3 palmitoylation.^32^ In cells incubated with Alk-4, DHHC7 overexpression increased IFITM3 labeling, while overexpression of a dominant negative DHHC7 mutant decreased IFITM3 labeling (Figure 2d). These results indicate that Alk-4 is metabolized into a long chain fatty acid that can be used as a substrate by DHHC palmitoyltransferases for protein palmitoylation.

### Labeling of IFITM3 by Alk-4 requires FASN

We have previously shown that IFITM3 palmitoylation is required for its antiviral activity against influenza virus infection.^22,33^ To determine if FASN-mediated *de novo* fatty acid biosynthesis contributes to an IFNβ-regulated IFITM3-mediated antiviral response, we measured endogenous IFITM3 labeling by Alk-4 in wild-type (WT) and FASN knockout HAP1 cells. As expected, IFNβ induced endogenous IFITM3 expression, and IFITM3 upregulation was independent of FASN expression (Figure 2e). In WT cells, Alk-4 treatment resulted in robust endogenous IFITM3 labeling that was absent in FASN-deficient cells (Figure 2f). Thus, we show for the first time that FASN contributes to the palmitoylation of endogenous IFITM3. Owing to the observations that IFITM3 is required for an effective IFNβ-mediated anti-influenza response,^32^ that palmitoylation of IFITM3 is required for its antiviral activity^32^, and that FASN regulates Alk-4 mediated IFITM3 palmitoylation (Figure 2f), we sought to determine the effect of FASN expression on IFNβ-mediated inhibition of influenza virus infection. In the absence of IFNβ, FASN expression had no effect on influenza infection (Figure 2g). However, IFNβ-mediated inhibition of influenza virus infection was significantly decreased in the absence of FASN expression (Figure 2g), suggesting that FASN-dependent palmitate synthesis likely contributes to the palmitoylation-dependent antiviral activity of IFITM3.

### Alk-4 labeling of myristoylated proteins is FASN dependent

Acetyl CoA is condensed with malonyl-CoA and elongated by FASN to generate palmitate for protein palmitoylation. To generate myristoyl CoA for myristoylation, palmitoyl CoA must be β-oxidized to myristoyl CoA before it is covalently attached to glycine residues by N-myristoyl transferases (Figure 4d). ^34,1^ To determine if Alk-4 is metabolized into a fatty acid analog that can selectively label myristoylated proteins, we tested whether a known myristoylated protein, HIV-1 matrix protein (MA), was labeled upon a 24-hour incubation with Alk-4. HEK293T cells were transfected with flag-tagged HIV-1 MA, or the myristoylation deficient matrix-G2A mutant (MA-G2A) and treated with Alk-4 or Alk-12 (an established chemical reporter of myristoylation). Immunoprecipitation of flag-tagged matrix and subsequent click reaction with azido-rhodamine revealed labeling of HIV-1 matrix in cells incubated with Alk-4 or Alk-12 (Figure 3a). The G2A-MA Gag protein, which cannot be myristoylated, was not labeled by Alk-4, indicating myristoylation site-specific labeling of HIV-1 matrix protein by Alk-4. Treatment of cells with Fasnall,^9,35^ a FASN inhibitor, abolished Alk-4 labeling of HIV-1 matrix protein. As a control, Fasnall treatment did not disrupt HIV-1 matrix protein labeling by Alk-12. To confirm the selective labeling of HIV-1 matrix protein that we observed with fluorescence-based click reactions, cell lysates were instead reacted with biotin azide. Biotin-conjugated proteins were precipitated with streptavidin agarose and bound proteins were released with sodium dithionite, which cleaves a diazo linker within the azido-biotin molecule, enabling selective elution of Alk-4 labeled proteins. Eluents were probed for the MA-Flag proteins, and, in the presence of Alk-4, HIV-1 matrix protein was recovered. HIV-1 matrix protein recovery was diminished both when a myristoylation deficient HIV-1 matrix protein variant was transfected (MA-G2A) (Figure 3b) and when FASN was inhibited by Fasnall. These results indicate that Alk-4 is metabolized into a fatty acid adduct that is specifically incorporated onto protein myristoylation sites in a FASN-dependent manner, and that can be detected by multiple labeling modalities.

**Figure 3:**
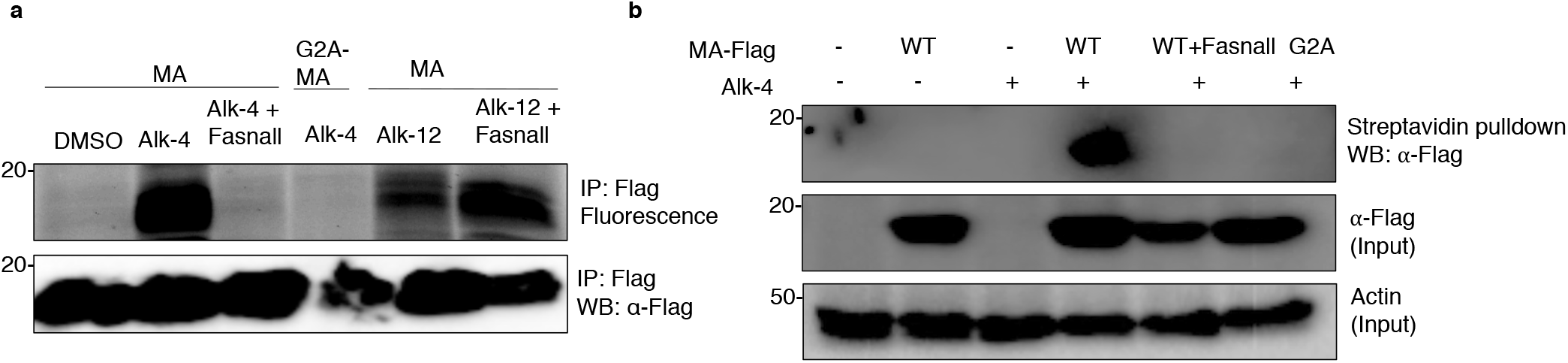
FASN-dependent incorporation of Alk-4 at known myristoylation sites. 293Ts were transfected overnight with plasmids encoding MA-Flag (MA), or the myristoylation deficient G2A-MA-Flag(G2A), and incubated for 24 hours either with Alk-4, or Alk-12. 10uM of the FASN inhibitor Fasnall was added one-hour post-transfection. **(a)** Immunoprecipitation of MA-Flag and click reaction with TAMRA azide revealed labeling of WT MA with Alk-4 and not G2A-MA, suggesting that Alk-4 labels MA at its known myristoylation site. FASN inhibition with Fasnall reduced labeling of MA by Alk-4, suggesting that FASN is required for labeling of MA by Alk-4. Alk-12 also labeled MA in both cells treated and untreated with Fasnall, indicating that Fasnall does not inhibit NMT function. **(b)** On click reaction with diazo azido biotin, streptavidin pulldown, and selective elution of Alk-4 labeled proteins, only WT-MA was labeled by Alk-4 and not G2A-MA, and Fasnall treatment also inhibited Alk-4 labeling of MA, corroborating our findings that FASN is required for labeling of MA by Alk-4 at its myristoylation site.

**Figure 4:**
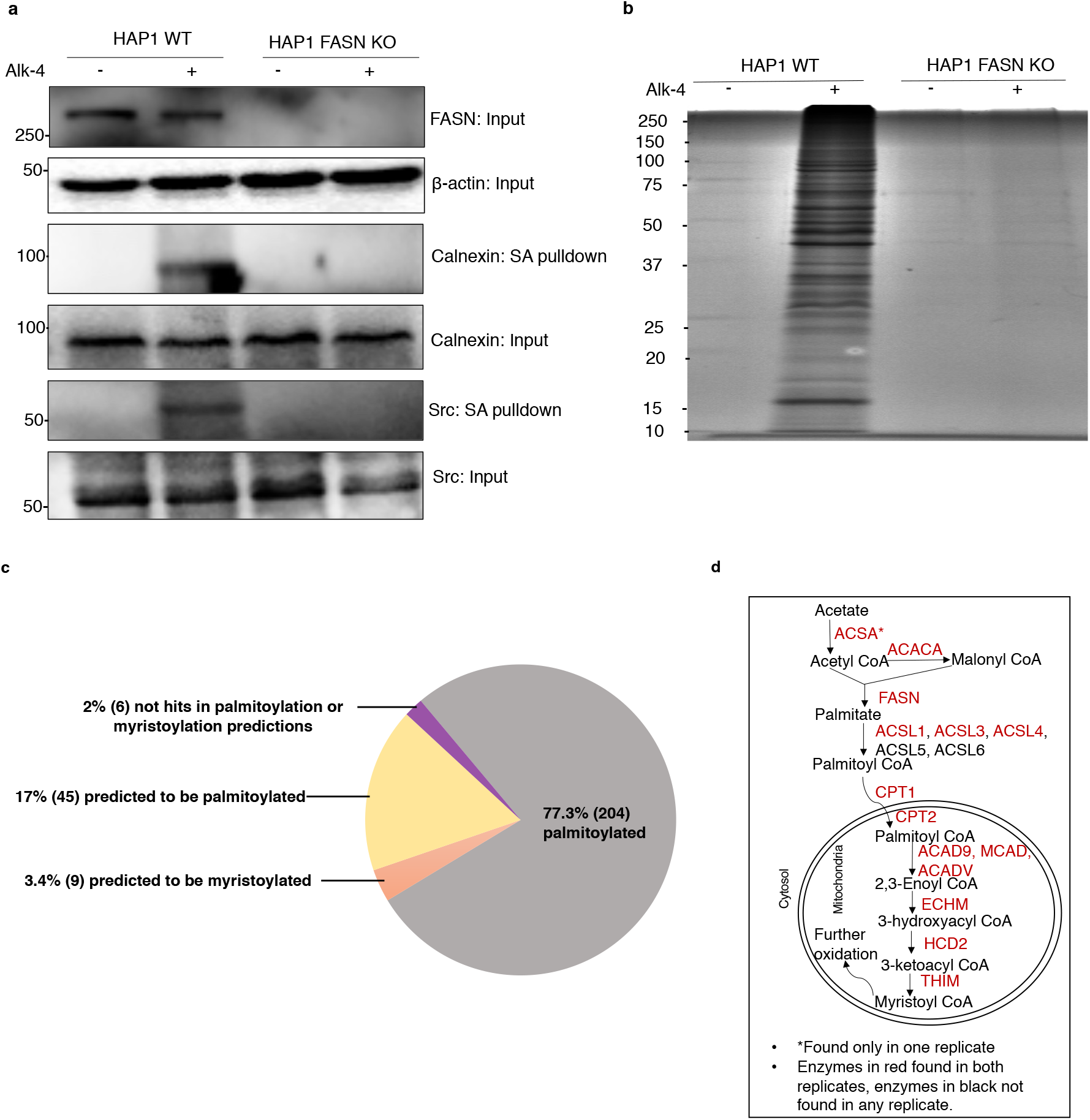
FASN-dependent purification of myristoylated and palmitoylated proteins from Alk-4 labeled cells. HAP1 wild-type (WT) and HAP1 FASN knockout (KO) cells were labeled with Alk-4, click reacted with diazo azido biotin, and labeled proteins were purified using streptavidin beads and eluted using sodium dithionite. **(a)** Western blotting indicates that proteins known to be palmitoylated (Calnexin) and myristoylated (Src) can be purified from cells in an Alk-4 and FASN dependent manner. **(b)** Coomassie staining of the streptavidin (SA) pulldown fraction revealed recovery of several proteins only in WT cells labeled with Alk-4. **(c)** Proteomics analysis of SA pulldown fraction revealed that 77% of proteins selectively recovered in the WT Alk-4 cells are found in at least one palmitoyl proteome or experimentally validated to be palmitoylated based on the SwissPalm database (Dataset 3). Additionally, 17% of proteins are predicted to be palmitoylated, based both on GPS-lipid analysis using high confidence settings, CSS-Palm, and SwissPalm (Dataset 1). GPS-Lipid analysis also revealed 3% of proteins are predicted to be myristoylated (consensus N-terminal Glycine and non-consensus sequence), while 2% of the proteins were not hits on prediction algorithms but had isoforms and cysteines as per SwissPalm analysis. **(d)** Proteomics analysis of SA pulldown fraction (n=2) revealed recovery of enzymes involved in elongation of short chain fatty acids, as well as enzymes involved in the mitochondrial beta-oxidation pathway for oxidation of long chain fatty acids. Long chain fatty acid such as palmitic acid and its oxidized product myristic acid can be used for fatty acylation of proteins.

### FASN-dependent, Alk-4 mediated metabolic labeling of endogenous fatty acylated proteins

To test the utility of Alk-4 as a global indicator of FASN-dependent protein acylation, we incubated the human fibroblast-like cell line HAP1 or a FASN-deficient clone of the HAP1 cells with Alk-4 or a vehicle control (DMSO). Following metabolic labeling of HAP1 cells with Alk-4, cell lysates were reacted with azido-biotin and labeled proteins were precipitated as described in Figure 3b. Eluents were then probed for proteins known to be palmitoylated (Calnexin)^36^ or myristoylated (Src).^37,38^ In wild-type HAP1 cells, Alk-4 labeling recovered both Calnexin and Src, while in FASN-deficient cells, incubation with Alk-4 did not enable Calnexin or Src recovery (Figure 4a). To determine the breadth of proteins recovered from cells incubated with Alk-4, we next used mass spectrometry to identify biotinylated proteins from HAP1 cells with or without Alk-4 and with or without FASN. FASN was only recovered from WT HAP1 cells incubated with Alk-4, consistent with the acyl intermediates formed between FASN and the elongating fatty acid chain (Figure 4d).^39^ This experiment also recovered, in an Alk-4 dependent manner, several enzymes involved in the metabolism of acetate to palmitate and myristate, including acetyl CoA synthetase (ACSA), acetyl CoA carboxylase 1 (ACACA), and multiple enzymes involved in fatty acid beta-oxidation (Figure 4d, Supplementary table 2). In total, Alk-4 labeling enabled recovery of 264 proteins in an Alk-4 and FASN-dependent manner (Figure 4b). Of these, 77% (203) have previously been identified in at least one palmitoyl proteome or they have been experimentally validated to be palmitoylated. These included well characterized palmitoylation substrates, such as Guanine nucleotide-binding protein G (GNAI1) and Catenin beta-1 (CTNB1) (Figure 4c, Supplementary table 1 & 3). Of the remaining proteins that were purified, 17% were predicted to be palmitoylated (e.g. SAM domain and HD domain-containing protein 1 [SAMHD1]), and 3% were predicted to be myristoylated (e.g. ribosomal protein S6 kinase alpha [KS6A1])^40,41^ (Figure 4c, Supplementary Table 1 & 4).

## Discussion

We describe a click-chemistry compatible FASN-substrate, Alk-4 (5-hexynoate), which selectively labels both palmitoylated (*e*.*g*. IFITM3, CD9) and myristoylated (*e*.*g*. HIV-1 matrix) proteins. Click chemistry compatible substrate analogs like Alk-4 overcome several inherent disadvantages of radiolabeling, such as long sample processing and film exposure times with low sensitivity.^16^ Moreover, click chemistry reactions can be combined with several detection methods that are compatible with high throughput applications including mass spectrometry-based proteomics, flow cytometry, fluorescence microscopy, and live cell imaging.^21^ Beyond identification of FASN-dependent protein acylation, Alk-4 has more functionality than a radiolabeled FASN substrate because it can be reacted with azido-biotin to facilitate streptavidin-based purification of FASN-dependent acylated proteins. We are not the first to suggest the utility of Alk-4, which has previously been evaluated as a chemical tool to monitor protein acetylation at shorter timescales, although it was noted that some of the acetylation reporters were incorporated onto proteins by chemical acylation.^24^ Nevertheless, our finding that FASN activity is required for Alk-4 labeling of multiple proteins such as IFITM3, HIV-1 matrix, Calnexin, and Src strongly supports the use of Alk-4 as a selective reporter of FASN-dependent protein acylation.

FASN activity is required for replication of several enveloped viruses, including Chikungunya,^10^ HIV-1,^9^ Influenza,^42^ and SARS-CoV-2,^43^ and many others.^6,7,13^ FASN inhibitors, including Fasnall^9,35^ and the TVB compounds^44,45^ have therapeutic potential, and pharmacological inhibition of FASN has been shown to modulate fatty acylation of viral proteins, including Chikungunya virus nsP1 palmitoylation,^46^ SARS-CoV-2 spike palmitoylation,^43^ and HIV-1 Gag myristoylation (Figure 3). In other cases, modulation of FASN activity affects host proteins that regulate infection, including MYD88 palmitoylation,^47^ and the results presented here that reveal that FASN activity is required for an effective IFNβ immune response against influenza virus, possibly by providing fatty acyl moieties for modification of IFITM3. Increased *de novo* FA biosynthesis and FASN up-regulation has also been observed in breast cancer, melanoma, and hepatocellular carcinoma^4^, and *de novo* fatty acid biosynthesis and lipogenesis has been shown to be essential for protein palmitoylation of Ras, Wnt^48^, Calnexin^49^, and Src^50^ in proliferating cells.^51,52^ In tumor cells, FASN inhibition can have consequences beyond inhibition of protein acylation^53^; *de novo* fatty acid biosynthesis has also been shown to be essential for membrane remodeling in tumor cells, where palmitate depletion via FASN inhibition led to disruption of lipid rafts and signaling pathways, ultimately resulting in apoptosis of tumor cells.^54^ Thus, *de novo* fatty acid biosynthesis is a broadly utilized, fundamental metabolic pathway exploited during carcinogenesis and virus replication, and Alk-4 and its ability to measure flux through the *de novo* FASN pathway provides a new tool to better understand the role of FASN-dependent protein acylation during FASN-dependent pathologies.

## METHODS

### Reagents, transfections, and infections

Reagents, including 5-hexynoic acid (Alk-4), were purchased from Sigma (St. Louis, MO) or Thermo Fisher Scientific, unless stated otherwise. Alk-12 and Alk-16 were synthesized by the Hang lab according to published protocols.^21^ Alk compounds were diluted in dimethyl sulfoxide (DMSO). HEK 293Ts were obtained from the American Type Culture Collection and were maintained in DMEM supplemented with 10% heat-inactivated fetal bovine serum (FBS; Serum Source International, Charlotte, NC) and 1% Penicillin/Streptomycin. HAP1 wild-type and HAP-1 FASN knockout cells were obtained from Horizon Discovery (Lafayette, Colorado) and were maintained in IMDM supplemented with 10% heat-inactivated FBS and 1% Penicillin/Streptomycin. HIV-1 matrix plasmids pQCXIP-MA-FH (pMA-Flag) and pQCXIP-G2A-FH (pG2A-MA-Flag) were kindly provided by Dr. Stephen Goff, Columbia University.^55^ HA-tagged IFITM3^22^ and HA-tagged ZDHHC7^56^ have been described. Cells were transfected using Genefect (Alkali Scientific, Fort Lauderdale, FL) or LipoJet transfection reagents (Signagen Laboratories, Rockville, MD), according to each manufacturer’s protocol. Fasnall was synthesized as described. ^35^ Influenza virus H1N1 strain PR8 was propagated in 10-day-old embryonated chicken eggs (Charles River) for 48 hours at 37 °C as described previously.^57,58^ IFNβ-treated HAP1 WT and FASN KO cells were treated with IFN overnight or left untreated as described previously^32^, and infected with H1N1 for 24 hours (MOI = 1). Cells were stained with anti-influenza virus nucleoprotein antibodies to measure percentage of infection by flow cytometry.

### Metabolic labeling, immunoprecipitations, and CuAAC

Cells were incubated for 24 hours with the indicated concentrations of Alk-3, Alk-4, Alk-12, Alk-16, or 0.001% DMSO in media supplemented with 1% charcoal-stripped FBS (Serum Source International, Catalog number FB02-500CS), and then collected and washed thrice in ice-cold 1x phosphate buffered saline (PBS). Cells were lysed for 10 minutes on ice in 100µl 1% Brij buffer (1% (w/v) Brij O10, 150mM NaCl, 50 mM triethanolamine with 1x EDTA-free complete protease inhibitor cocktail. Protein concentration was determined using the BCA assay. Flag precipitations used 500 µg of cell lysate mixed with Protein G coated agarose beads and incubated with Anti-Flag antibody (catalog number F3165) for 2 hours at 4°C.Anti-HA IPs were performed using EZview Red Anti-HA Affinity Gel. Protein-conjugated beads were washed thrice with radioimmunoprecipitation assay (RIPA) buffer (50 mM triethanolamine, 150 mM NaCl, 1% sodium deoxycholate, 1% triton-X-100, 0.1 % SDS). Protein complexes bound to antibody coated beads were released by adding 4% SDS buffer (150 mM NaCl, 50 mM triethanolamine, 4% [w/v] SDS), and the click reaction was initiation by addition of 2.75µl of click chemistry master mix (0.5µl of 5mM azido-rhodamine or tetramethylrhodamine-5-carbonyl azide [Click Chemistry Tools, Scottsdale, AZ] in DMSO, 0.5µl of 50mM tris(2-carboxyethyl)phosphine [TCEP], 0.5µl of 50mM CuSO4, and 1.5µl of 2mM tris (1-benzyl-1H-1,2,3-triazol-4-yl)methyl)amine (TBTA) in 1:4 [v/v] DMSO/butanol). Reactions were incubated for one hour at room temperature, and proteins were eluted from the beads by heating at 95°C for 5 minutes in 4X SDS sample loading buffer (40% (v/v) glycerol, 240 mM Tris·Cl, pH 6.8, 8% (w/v) sodium dodecyl sulfate (SDS), 0.04% (w/v) bromophenol blue, 5% 2-mercaptoethanol). Eluted proteins were resolved on 4-20% tris-glycine gels. To detect fluorescently labeled proteins, the gel was destained in 40% distilled water(v/v), 50%(v/v) methanol, 10%(v/v) acetic acid and visualized using on an Amersham Typhoon 9410 with 532-nm excitation and 580-nm detection filters. For biotin-based click reactions, Alk-4 incubated cells were lysed with 50µl 4% SDS buffer with 1x EDTA-free protease inhibitors supplemented with benzonase nuclease (Catalog number E1014). One mg of cell lysate was resuspended in 445µl 1x SDS buffer with 1x EDTA-free protease inhibitors and incubated for 1.5 hours with 55µl of click reaction master-mix consisting of 10µl of 5mM diazo biotin azide (Click Chemistry Tools), 10µl of 50mM TCEP, 25 ul of 2mM TBTA, 10µl of 50mM CuSO4. Proteins were precipitated using chloroform-methanol to remove unreacted biotin azide, and the precipitant was resuspended in 100µl 4% SDS buffer containing protease inhibitors supplemented and 2µl of 0.5M EDTA solution to chelate residual copper. Equivalent amount of protein in 100µl 4%SDS buffer and 200 µl 1% Brij buffer with EDTA-free protease inhibitors was incubated with 75ul streptavidin agarose (EMD Millipore) for two hours at room temperature. Protein-conjugated beads were washed once in PBS/0.2-1% SDS, and thrice in PBS. Labeled proteins were selectively eluted by two elutions with 50mM sodium dithionite, desalted using spin desalting columns, mixed with 4X SDS sample loading buffer, and resolved on 10-12% Tris-Glycine gel. Coomassie staining was done using standard techniques.^59^

### Western blotting

Proteins were transferred onto PVDF membrane (Bio-Rad, Hercules, CA), blocked with 5% bovine serum albumin dissolved in 1x tris-buffered saline (TBS) containing 0.1% Tween-20. Anti-Flag (catalog number F3165) and Anti-FASN (catalog number SAB4300700) antibodies were purchased from Sigma and used at a final concentration of 1:1000; anti-Calnexin (catalog number 2679S) and anti-Src (catalog number 2109S) antibodies were purchased from Cell Signaling Technologies (Danvers, MA) and used at a final concentration of 1:2000. Anti-rabbit secondary antibody (Thermo Fisher Scientific catalog number 31460) was used at a final concentration of 1: 5000, and anti-mouse secondary antibody (Cell Signaling Technology catalog number 7076S) was used at a final concentration of 1:2000.

### Protein identification

Capillary-LC-MS/MS was performed using a Thermo Scientific orbitrap fusion mass spectrometer equipped with an EASY-Spray™ Sources operated in positive ion mode. Samples were separated on an easy spray nano column (Pepmap™ RSLC, C18 3µ 100A, 75µm X150mm Thermo Scientific) using a 2D RSLC HPLC system from Thermo Scientific. The full scan was performed at FT mode and the resolution was set at 120,000. EASY-IC was used for internal mass calibration. Mass spectra were searched using Mascot Daemon by Matrix Science version 2.3.2 (Boston, MA) and the database searched against Uniprot Human database (version 12032015). Data from two independent experiments were compiled on Scaffold Visualization software (Scaffold 4.9.0, Proteome Software Inc., Portland, OR). Proteins were identified based on total spectrum count with a 1% false discovery rate (FDR) and a minimum of two peptides. Proteins were considered high confidence hits if they had DMSO spectral count of zero, and a minimum spectral count of five in both replicates. Putative fatty acylation sites in high confidence proteins were identified in the SwissPalm protein S-palmitoylation database (Version 3, https://SwissPalm.org/) using Dataset 3 (proteins found in at least one palmitoyl-proteome or experimentally validated to be palmitoylated) and Dataset 1 (All datasets). Protein sequences were also searched against GPS-Lipid using high threshold settings (Version 1.0, http://lipid.biocuckoo.org/), as described in the supplementary methods section.

## Acknowledgements

This work was supported by NIH/NIAID grants AI141037 to J.J.K and AI130110 and AI142256 to J.S.Y., and The Ohio State University Department of Microbiology. We thank Dr. Howard Hang of The Scripps Research Institute for providing Alk-12, Alk-16, and azido-rhodamine, and Dr. Yiping Zhu and Dr. Stephen Goff of Columbia University for providing MA-Flag and G2A-MA-Flag constructs. We thank Dr. Liwen Zhang of the OSU Mass spectrometry and Proteomics facility for acquisition and processing of the LC-MS/MS data under support from NIH Grants CA016058 and OD018056.

## Author Contributions

K.P.K., J.S.Y., and J.J.K. conceived the project and wrote the paper. K.P.K. and L.Z. performed all experiments. K.P.K., L.Z., J.S.Y., and J.J.K. performed data analysis. M.B., D.R.L., and T.A.J.H. provided experimental expertise. All authors edited and approved the manuscript.

## References

1. Resh, M. M. Fatty acylation of proteins: The long and the short of it. Prog Lipid Res 63, 120–131 (2016).

2. Suburu, J. et al.. Fatty acid metabolism: Implications for diet, genetic variation, and disease. Food Bioscience (2013). doi:10.1016/j.fbio.2013.07.003

3. Liu, H., Liu, J. Y., Wu, X. & Zhang, J. J. Biochemistry, molecular biology, and pharmacology of fatty acid synthase, an emerging therapeutic target and diagnosis/prognosis marker. International Journal of Biochemistry and Molecular Biology (2010).

4. Kuhajda, F. F. Fatty-acid synthase and human cancer: New perspectives on its role in tumor biology. Nutrition (2000). doi:10.1016/S0899-9007(99)00266-X

5. Zhang, H. et al.. Differential regulation of host genes including hepatic fatty acid synthase in HBV-transgenic mice. J Proteome Res 12, 2967–2979 (2013).

6. Tongluan, N. et al.. Involvement of fatty acid synthase in dengue virus infection. Virol J 14, 28 (2017).

7. Li, Y., Webster-Cyriaque, J., Tomlinson, C. C., Yohe, M. & Kenney, S. Fatty Acid Synthase Expression Is Induced by the Epstein-Barr Virus Immediate-Early Protein BRLF1 and Is Required for Lytic Viral Gene Expression. J. Virol. (2004). doi:10.1128/jvi.78.8.4197-4206.2004

8. Nasheri, N. et al.. Modulation of fatty acid synthase enzyme activity and expression during hepatitis C virus replication. Chem. Biol. 20, 570–82 (2013).

9. Kulkarni, M. M. et al.. Cellular fatty acid synthase is required for late stages of HIV-1 replication. Retrovirology 14, 45 (2017).

10. Bakhache, W. et al.. Fatty acid synthase and stearoyl-CoA desaturase-1 are conserved druggable cofactors of Old World Alphavirus genome replication. Antiviral Res. (2019). doi:10.1016/j.antiviral.2019.104642

11. Zhang, N., Zhao, H. & Zhang, L. Fatty Acid Synthase Promotes the Palmitoylation of Chikungunya Virus nsP1. J. Virol. (2018). doi:10.1128/jvi.01747-18

12. Krishnan, M. N. et al.. RNA interference screen for human genes associated with West Nile virus infection. Nature 455, 242–245 (2008).

13. Martín-Acebes, M. A., Blázquez, A. B., Jiménez de Oya, N., Escribano-Romero, E. & Saiz, J. J. West Nile virus replication requires fatty acid synthesis but is independent on phosphatidylinositol-4-phosphate lipids. PLoS One 6, e24970 (2011).

14. Hang, H. H. & Linder, M. E. Exploring protein lipidation with chemical biology. Chemical Reviews (2011). doi:10.1021/cr2001977

15. Rodenburg, R. N. P. et al.. Stochastic palmitoylation of accessible cysteines in membrane proteins revealed by native mass spectrometry. Nat. Commun. (2017). doi:10.1038/s41467-017-01461-z

16. Draper, J. M. & Smith, C. C. Palmitoyl acyltransferase assays and inhibitors (Review).Molecular Membrane Biology (2009). doi:10.1080/09687680802683839

17. Gao, X. & Hannoush, R. R. A Decade of Click Chemistry in Protein Palmitoylation: Impact on Discovery and New Biology. Cell Chemical Biology (2018). doi:10.1016/j.chembiol.2017.12.002

18. Yap, M. C. et al.. Rapid and selective detection of fatty acylated proteins using ω-alkynyl-fatty acids and click chemistry. J. Lipid Res. (2010). doi:10.1194/jlr.D002790

19. Thiele, C. et al.. Tracing fatty acid metabolism by click chemistry. ACS Chem Biol 7, 2004–2011 (2012).

20. Ourailidou, M. E., Zwinderman, M. R. H. & Dekker, F. F. Bioorthogonal metabolic labelling with acyl-CoA reporters: Targeting protein acylation. MedChemComm (2016). doi:10.1039/c5md00446b

21. Charron, G. et al.. Robust fluorescent detection of protein fatty-acylation with chemical reporters. J Am Chem Soc 131, 4967–4975 (2009).

22. Yount, J. S. et al.. Palmitoylome profiling reveals S-palmitoylation-dependent antiviral activity of IFITM3. Nat Chem Biol 6, 610–614 (2010).

23. Chesarino, N. M. et al.. Chemoproteomics reveals Toll-like receptor fatty acylation. BMC Biol 12, 91 (2014).

24. Yang, Y. Y., Ascano, J. M. & Hang, H. H. Bioorthogonal chemical reporters for monitoring protein acetylation. J Am Chem Soc 132, 3640–3641 (2010).

25. Mitchell, D. A., Vasudevan, A., Linder, M. E. & Deschenes, R. R. Protein palmitoylation by a family of DHHC protein S-acyltransferases. Journal of Lipid Research (2006). doi:10.1194/jlr.R600007-JLR200

26. Greaves, J. et al.. Molecular basis of fatty acid selectivity in the zDHHC family of S- acyltransferases revealed by click chemistry. Proc Natl Acad Sci U S A 114, E1365– E1374 (2017).

27. Jennings, B. C. & Linder, M. M. DHHC protein S-acyltransferases use similar ping-pong kinetic mechanisms but display different acyl-CoA specificities. J Biol Chem 287, 7236–7245 (2012).

28. Muszbek, L., Haramura, G., Cluette-Brown J. E., Van Cott E. M. & Laposata, M. The pool of fatty acids covalently bound to platelet proteins by thioester linkages can be altered by exogenously supplied fatty acids. Lipids 34 Suppl, S331–7 (1999).

29. Hallak, H. et al.. Covalent binding of arachidonate to G protein alpha subunits of human platelets. J Biol Chem 269, 4713–4716 (1994).

30. Veit, M., Reverey, H. & Schmidt, M. M. Cytoplasmic tail length influences fatty acid selection for acylation of viral glycoproteins. Biochem J 318 (Pt 1, 163–172 (1996).

31. Thinon, E., Fernandez, J. P., Molina, H. & Hang, H. H. Selective Enrichment and Direct Analysis of Protein S-Palmitoylation Sites. J Proteome Res 17, 1907–1922 (2018).

32. McMichael, T. M. et al.. The palmitoyltransferase ZDHHC20 enhances interferon-induced transmembrane protein 3 (IFITM3) palmitoylation and antiviral activity. J Biol Chem 292, 21517–21526 (2017).

33. Percher, A. et al.. Mass-tag labeling reveals site-specific and endogenous levels of protein S-fatty acylation. Proc Natl Acad Sci U S A 113, 4302–4307 (2016).

34. Farazi, T. A., Waksman, G. & Gordon, J. J. The biology and enzymology of protein N- myristoylation. J Biol Chem 276, 39501–39504 (2001).

35. Alwarawrah, Y. et al.. Fasnall, a Selective FASN Inhibitor, Shows Potent Anti-tumor Activity in the MMTV-Neu Model of HER2(+) Breast Cancer. Cell Chem Biol 23, 678–688 (2016).

36. Lakkaraju, A. K. K. et al.. Palmitoylated calnexin is a key component of the ribosome-translocon complex. EMBO J. (2012). doi:10.1038/emboj.2012.15

37. Patwardhan, P. & Resh, M. M. Myristoylation and Membrane Binding Regulate c-Src Stability and Kinase Activity. Mol. Cell. Biol. (2010). doi:10.1128/mcb.00246-10

38. Resh, M. M. Myristylation and palmitylation of Src family members: The fats of the matter. Cell (1994). doi:10.1016/0092-8674(94)90104-X

39. Wakil, S. S. Fatty acid synthase, a proficient multifunctional enzyme. Biochemistry 28, 4523–4530 (1989).

40. Xie, Y. et al.. GPS-Lipid: A robust tool for the prediction of multiple lipid modification sites. Sci. Rep. (2016). doi:10.1038/srep28249

41. Ren, J. et al.. CSS-Palm 2.0: An updated software for palmitoylation sites prediction. Protein Eng. Des. Sel. (2008). doi:10.1093/protein/gzn039

42. Munger, J. et al.. Systems-level metabolic flux profiling identifies fatty acid synthesis as a target for antiviral therapy NIH Public Access Author Manuscript. Nat Biotechnol 26, 1179–1186 (2008).

43. Lee, M. et al.. Fatty acid synthase inhibition prevents palmitoylation of SARS-CoV2 spike protein and improves survival of mice infected with murine hepatitis virus. Preprint at https://www.biorxiv.org/content/10.1101/2020.12.20.423603v1 (2020). doi:10.1101/2020.12.20.423603

44. Aquino, I. I. de et al. Anticancer properties of the fatty acid synthase inhibitor TVB-3166 on oral squamous cell carcinoma cell lines. Arch. Oral Biol. (2020). doi:10.1016/j.archoralbio.2020.104707

45. Zaytseva, Y. Y. et al.. Preclinical evaluation of novel fatty acid synthase inhibitors in primary colorectal cancer cells and a patient-derived xenograft model of colorectal cancer. Oncotarget 9, 24787–24800 (2018).

46. Zhang, N., Zhao, H. & Zhang, L. Fatty Acid Synthase Promotes the Palmitoylation of Chikungunya Virus nsP1. J Virol 93, (2019).

47. Kim, Y.-C. et al.. Toll-like receptor mediated inflammation requires FASN-dependent MYD88 palmitoylation. Nat. Chem. Biol. (2019). doi:10.1038/s41589-019-0344-0

48. Kaemmerer, E. & Gassler, N. Wnt lipidation and modifiers in intestinal carcinogenesis and cancer. Cancers (2016). doi:10.3390/cancers8070069

49. Lynes, E. M. et al.. Palmitoylation is the switch that assigns calnexin to quality control or ER Ca2+ signaling. J. Cell Sci. (2013). doi:10.1242/jcs.125856

50. Kim, S. et al.. Blocking myristoylation of Src inhibits its kinase activity and suppresses prostate cancer progression. Cancer Res. (2017). doi:10.1158/0008-5472.CAN-17-0981

51. Fiorentino, M. et al.. Overexpression of fatty acid synthase is associated with palmitoylation of Wnt1 and cytoplasmic stabilization of beta-catenin in prostate cancer. Lab Invest 88, 1340–1348 (2008).

52. Yao, C. H. et al.. Exogenous Fatty Acids Are the Preferred Source of Membrane Lipids in Proliferating Fibroblasts. Cell Chem Biol 23, 483–493 (2016).

53. Chen, M. & Huang, J. The expanded role of fatty acid metabolism in cancer: new aspects and targets. Precis. Clin. Med. 2, 183–191 (2019).

54. Ventura, R. et al.. Inhibition of de novo Palmitate Synthesis by Fatty Acid Synthase Induces Apoptosis in Tumor Cells by Remodeling Cell Membranes, Inhibiting Signaling Pathways, and Reprogramming Gene Expression. EBIOM 2, 808–824 (2015).

55. Zhu, Y. et al.. Heme Oxygenase 2 Binds Myristate to Regulate Retrovirus Assembly and TLR4 Signaling. Cell Host Microbe (2017). doi:10.1016/j.chom.2017.01.002

56. Hach, J. C., McMichael, T., Chesarino, N. M. & Yount, J. J. Palmitoylation on Conserved and Nonconserved Cysteines of Murine IFITM1 Regulates Its Stability and Anti-Influenza A Virus Activity. J. Virol. 87, 9923–9927 (2013).

57. Yount, J. S., Kraus, T. A., Horvath, C. M., Moran, T. M. & López, C. C. A Novel Role for Viral-Defective Interfering Particles in Enhancing Dendritic Cell Maturation. J. Immunol. (2006). doi:10.4049/jimmunol.177.7.4503

58. Moltedo, B., Li, W., Yount, J. S. & Moran, T. T. Unique type I interferon responses determine the functional fate of migratory lung dendritic cells during influenza virus infection. PLoS Pathog. (2011). doi:10.1371/journal.ppat.1002345

59. Merril, C. C. Gel-staining techniques. Methods Enzymol. (1990). doi:10.1016/0076-6879(90)82038-4

