## Supplementary material for "A bioorthogonal chemical reporter for fatty acid synthase-dependent protein acylation"

**Supplementary methods:**

**Protein identification using Capillary-LC-MS/MS:**

Samples were resuspended in 50mM ammonium bicarbonate (ABC) solution containing DTT and incubated at 65°C for 15 min followed by addition of 50 mM ABC and 15 $\mu$g/$\mu$L iodoacetamide. Sequencing grade trypsin dissolved in 50mM ABC was added and digested overnight at 37°C. Peptides were acidified by adding 0.1% Formic acid. The sample was dried in a vacufuge and resuspended in 50 mM acetic acid. Capillary-liquid chromatography-nanospray tandem mass spectrometry (Capillary-LC/MS/MS) for protein identification was performed on a Thermo Scientific orbitrap Fusion mass spectrometer equipped with an EASY-Spray™ Sources operated in positive ion mode. Samples were separated on an easy spray nano column (Pepmap$^{TM}$ RSLC, C18 3$\mu$ 100A, 75$\mu$m X150mm Thermo Scientific) using a 2D RSLC HPLC system from Thermo Scientific. Each sample was injected into the $\mu$-Precolumn Cartridge (Thermo Scientific,) and desalted with 0.1% Formic Acid in water for 5 minutes. Mobile phase A was 0.1% Formic Acid in water and acetonitrile (with 0.1% formic acid) was used as mobile phase B. Flow rate was set at 300nL/min. Mobile phase A was 0.1% Formic Acid in water and acetonitrile (with 0.1% formic acid) was used as mobile phase B. Flow rate was set at 300nL/min. Typically, mobile phase B was kept at 2% for 5 min and then increased from 2% to 16% in 100 min and then increased from 16 to 25% in 20min and again from 55%-85% in 1 min and then kept at 85% for another 4 min before being brought back quickly to 2% in 1 min.MS/MS data was acquired with a spray voltage of 1.6 KV and a capillary temperature of 305°C. The full scan was performed at FT mode and the resolution was set at 120,000. EASY-IC was used for internal mass calibration. Mass spectra were searched using Mascot Daemon by Matrix Science version 2.3.2 (Boston, MA) and the database searched against Uniprot Human database (version 12032015). The mass accuracy of the precursor ions were set to 10ppm, accidental pick of 1 13C peaks was also included into the search. The fragment mass tolerance was set to 0.5 Da. Considered variable modifications were oxidation (Met), deamidation (N and Q) and carbamidomethylation (Cys). Four missed cleavages for the enzyme were permitted. A decoy database was also searched to determine the false discovery rate (FDR) and peptides were filtered according to the FDR. The significance threshold was set at $p<0.05$ and bold red peptides is required for valid peptide identification. Proteins with less than 1% FDR as well as a minimal of 2 significant peptides detected were considered as valid proteins.

Data from two independent experiments were compiled on Scaffold Visualization software (Version 4, Proteome Software, https://www.proteomesoftware.com/). Proteins were identified based on total spectrum count with set conditions of 1% false discovery rate (FDR) and 2 minimum number of peptides. Proteins were considered as high confidence hits if they had DMSO spectral count=0, and minimum spectral count of at least 5 in both replicates. The 264 high confidence proteins were searched against the SWISSPALM protein S-palmitoylation database (Version 3, https://swisspalm.org/) using Dataset 3 (proteins found in at least 1 palmitoyl-proteome or experimentally validated to be palmitoylated) and Dataset 1 (All datasets). Protein sequences were also searched against GP

S-Lipid using high threshold settings (Version 1.0, http://lipid.biocuckoo.org/).

**Supplementary table 1: Proteomic analysis of HAP1 wild-type (WT) and FASN knockout (KO) cells reveal selective recovery in WT alk-4 labeled cells of several proteins identified in other palmitoyl proteome experiments based on the database SwissPalm, as well as other predicted fatty acyl modifications (n=2).** Sequences were visualized and retrieved using Scaffold viewer (number of minimum peptides=2, 1% false discovery rate). For each replicate, proteins were considered as “high confidence” hits based on total spectral count if they had DMSO spectral count =0 and minimum spectral count=5. For the “pooled” set, only proteins which met the above criteria in both the replicates were chosen as high confidence candidates. No proteins were identified in the KO alk-4 “pooled” set. SwissPalm (Dataset 3) was used to identify proteins whose palmitoylation has been experimentally validated or have been found in at least one palmitoyl proteome study. Proteins were also analyzed using Dataset 1 in SwissPalm, which includes the criteria in Dataset 3 and also considers number of cysteines, orthologs, and palmitoylation sites of a protein. GPS-lipid analysis was done using high confidence settings; for myristoylation, sequence analysis was used to count the number of N-terminal glycines.

| **Modification** | **Palmitoylation** | | | **Myristoylation** | |
|---|---|---|---|---|---|
| Technique or Database | SwissPalm Dataset 3 (number of proteins found in palmitoyl proteomes or experimentally validated) | SwissPalm Dataset 1 (number of proteins found in palmitoyl proteomes or experimentally validated, or number of predicted palmitoylated sites or cysteines or orthologs) | GPS -Lipid prediction | GPS-Lipid prediction | Number of proteins with N-terminal Glycines |
| Replicate 1 (589 proteins) | 433 (73%) | 588 (99.8%) | 350 (59.4%) | 123 (20.9%) | 39 (6.6%) |
| Replicate 2 (526 proteins) | 387 (73%) | 524 (99.6%) | 248 (47.1%) | 65 (12.4%) | 27 (5.1%) |
| Pooled (264 proteins) | 203 (76%) | 264 (100%) | 133 (50.3%) | 33 (12.5%) | 14 (5.3%) |

**Supplementary table 2: Proteomics analysis of streptavidin pulldown fraction (n=2) revealed recovery of enzymes involved in elongation of short chain fatty acids with Alk-4, as well as enzymes involved in the mitochondrial beta-oxidation pathway for oxidation of long chain fatty acids.** Long chain fatty acid such as palmitic acid and its oxidized product myristic acid can be used for fatty acylation of proteins, and streptavidin pulldown and proteomics analysis of Alk-4 labeled cells revealed recovery of the enzymes involved in elongation as well as oxidation of fatty acids, as revealed by the total spectral counts below.

| Enzyme | Total spectral count (Replicate 1, Replicate 2) | | | |
|---|---|---|---|---|
| | WT DMSO | WT ALK-4 | KO DMSO | KO Alk-4 |
| Acetyl CoA synthetase (ACSA) | 0, - | 3, - | 0, - | 0, - |
| Acyl CoA carboxylase1 (ACACA) | 0, 1 | 21, 13 | 2, 0 | 5, 1 |
| Fatty acid synthase(FASN) | 3, 35 | 124, 165 | 0, 0 | 0, 0 |
| Long chain fatty acid ligase 1 (ACSL1) | 0 | 6, 14 | 0 | 2 |
| Long chain fatty acid ligase 1 (ACSL3) | 0, 0 | 12, 10 | 0, 0 | 2, 0 |
| Long chain fatty acid CoA ligase 4 (ACSL4) | 0, 0 | 7, 11 | 0, 0 | 0, 0 |
| Carnitine-O-palmitoyltransferase-1 (CPT1) | 0, 0 | 8, 9 | 0, 0 | 0, 0 |
| Carnitine-O-palmitoyltransferase-2 (CPT2) | 0, 0 | 6, 15 | 1, 0 | 1, 0 |
| Acyl-CoA dehydrogenase 9 (ACAD9) | 0, 0 | 5, 7 | 0, 0 | 1, 0 |
| Medium-chain specific acyl-CoA dehydrogenase (MCAD) | 0, 0 | 7, 3 | 0, 0 | 2, 0 |
| Very long-chain specific acyl CoA dehydrogenase (ACADV) | 0, 0 | 5, 10 | 0, 0 | 0, 0 |
| Enoyl-CoA hydratase (ECHM) | 0, 3 | 11, 16 | 1, 0 | 6, 1 |
| 3-hydroxyacyl-CoA dehydrogenase (HCD2) | 0, 6 | 13, 25 | 0, 0 | 4, 5 |
| 3-ketoacyl thiolase (THIM) | 0, 4 | 13, 24 | 0, 0 | 5, 7 |

**Supplementary table 3:** Out of 264 proteins recovered in WT HAP1 cells labeled with Alk-4 using high confidence settings as described in Figure 4 and Supplementary table, 204 (77%) of proteins have been found in palmitoyl proteomes or experimentally validated to be palmitoylated ("Techniques") using Dataset 3 of the SWISSPALM database.

| Query Identifier | Found in palmitoyl-proteomes | Techniques | Times validated |
|---|---|---|---|
| MPRI_HUMAN | 7/17 | 4 | 1 |
| RAB8A_HUMAN | 7/17 | 4 | 0 |
| NDKB_HUMAN | 6/17 | 4 | 0 |
| CTND1_HUMAN | 6/17 | 3 | 2 |
| ATPG_HUMAN | 6/17 | 3 | 0 |
| CPT1A_HUMAN | 6/17 | 3 | 0 |
| GNAI1_HUMAN | 5/17 | 3 | 7 |
| CTNB1_HUMAN | 5/17 | 3 | 4 |
| LETM1_HUMAN | 5/17 | 3 | 0 |
| NSUN2_HUMAN | 5/17 | 3 | 0 |
| TBB6_HUMAN | 5/17 | 3 | 0 |
| DSRAD_HUMAN | 5/17 | 3 | 0 |
| RAB14_HUMAN | 5/17 | 2 | 1 |
| ACSL4_HUMAN | 4/17 | 3 | 0 |
| DDX46_HUMAN | 4/17 | 3 | 0 |
| TMEDA_HUMAN | 4/17 | 3 | 0 |
| RRP12_HUMAN | 4/17 | 3 | 0 |
| DHB12_HUMAN | 4/17 | 3 | 0 |
| CHD4_HUMAN | 4/17 | 3 | 0 |
| ACSL3_HUMAN | 4/17 | 3 | 0 |
| SRPRB_HUMAN | 4/17 | 3 | 0 |
| ADT1_HUMAN | 4/17 | 3 | 0 |
| FDFT_HUMAN | 4/17 | 3 | 0 |
| 1C17_HUMAN | 4/17 | 3 | 0 |
| AL3A2_HUMAN | 4/17 | 3 | 0 |
| ACAD9_HUMAN | 4/17 | 3 | 0 |
| AP2A2_HUMAN | 4/17 | 3 | 0 |
| CN37_HUMAN | 4/17 | 3 | 0 |

| Query Identifier | Found in palmitoyl-proteomes | Techniques | Times validated |
|---|---|---|---|
| PLAK_HUMAN | 4/17 | 2 | 1 |
| APMAP_HUMAN | 4/17 | 1 | 0 |
| RU17_HUMAN | 3/17 | 3 | 0 |
| RECQ1_HUMAN | 3/17 | 3 | 0 |
| ERP29_HUMAN | 3/17 | 3 | 0 |
| VAPB_HUMAN | 3/17 | 3 | 0 |
| NCBP1_HUMAN | 3/17 | 3 | 0 |
| PTN1_HUMAN | 3/17 | 3 | 0 |
| DRG1_HUMAN | 3/17 | 3 | 0 |
| ZCCHV_HUMAN | 3/17 | 3 | 0 |
| IF2B_HUMAN | 3/17 | 3 | 0 |
| TNPO3_HUMAN | 3/17 | 3 | 0 |
| TBL3_HUMAN | 3/17 | 3 | 0 |
| GNL3_HUMAN | 3/17 | 3 | 0 |
| NAT10_HUMAN | 3/17 | 3 | 0 |
| ADAS_HUMAN | 3/17 | 3 | 0 |
| MK01_HUMAN | 3/17 | 3 | 0 |
| SPTC1_HUMAN | 3/17 | 3 | 0 |
| VATB2_HUMAN | 3/17 | 3 | 0 |
| NU205_HUMAN | 3/17 | 3 | 0 |
| PGAM5_HUMAN | 3/17 | 3 | 0 |
| SNUT2_HUMAN | 3/17 | 3 | 0 |
| UCRI_HUMAN | 3/17 | 2 | 0 |
| PRP6_HUMAN | 3/17 | 2 | 0 |
| DCTN1_HUMAN | 3/17 | 2 | 0 |
| RAB8B_HUMAN | 3/17 | 2 | 0 |
| AL1B1_HUMAN | 3/17 | 2 | 0 |

| UniProt ID | Found in palmitoyl-proteomes | Techniques | Times validated |
|---|---|---|---|
| MYO1C_HUMAN | 3/17 | 2 | 0 |
| CYFP1_HUMAN | 3/17 | 2 | 0 |
| NCLN_HUMAN | 3/17 | 2 | 0 |
| SYIM_HUMAN | 3/17 | 1 | 0 |
| GEMI5_HUMAN | 2/17 | 4 | 0 |
| DKC1_HUMAN | 2/17 | 3 | 0 |
| ACINU_HUMAN | 2/17 | 3 | 0 |
| WDR3_HUMAN | 2/17 | 3 | 0 |
| WDR12_HUMAN | 2/17 | 3 | 0 |
| ADK_HUMAN | 2/17 | 3 | 0 |
| AP3D1_HUMAN | 2/17 | 3 | 0 |
| BAZ1B_HUMAN | 2/17 | 3 | 0 |
| TPP2_HUMAN | 2/17 | 3 | 0 |
| BYST_HUMAN | 2/17 | 3 | 0 |
| SUCB1_HUMAN | 2/17 | 3 | 0 |
| STAT1_HUMAN | 2/17 | 3 | 0 |
| SRP54_HUMAN | 2/17 | 3 | 0 |
| CMC2_HUMAN | 2/17 | 3 | 0 |
| ABCF1_HUMAN | 2/17 | 3 | 0 |
| SPSY_HUMAN | 2/17 | 3 | 0 |
| RIF1_HUMAN | 2/17 | 3 | 0 |
| DDX56_HUMAN | 2/17 | 3 | 0 |
| RBM14_HUMAN | 2/17 | 3 | 0 |
| DHX30_HUMAN | 2/17 | 3 | 0 |
| DYL1_HUMAN | 2/17 | 3 | 0 |
| EIF2A_HUMAN | 2/17 | 3 | 0 |
| ELOC_HUMAN | 2/17 | 3 | 0 |
| POP1_HUMAN | 2/17 | 3 | 0 |

| UniProt ID | Found in palmitoyl-proteomes | Techniques | Times validated |
|---|---|---|---|
| FERM2_HUMAN | 2/17 | 3 | 0 |
| PESC_HUMAN | 2/17 | 3 | 0 |
| GBB2_HUMAN | 2/17 | 3 | 0 |
| GLSK_HUMAN | 2/17 | 3 | 0 |
| HEAT1_HUMAN | 2/17 | 3 | 0 |
| HMGB2_HUMAN | 2/17 | 3 | 0 |
| HNRL2_HUMAN | 2/17 | 3 | 0 |
| TTL12_HUMAN | 2/17 | 2 | 0 |
| NCKP1_HUMAN | 2/17 | 2 | 0 |
| RBGPR_HUMAN | 2/17 | 2 | 0 |
| ACSL1_HUMAN | 2/17 | 2 | 0 |
| PSIP1_HUMAN | 2/17 | 2 | 0 |
| EDC4_HUMAN | 2/17 | 2 | 0 |
| DDX50_HUMAN | 2/17 | 2 | 0 |
| HDAC1_HUMAN | 2/17 | 2 | 0 |
| PDLI5_HUMAN | 2/17 | 2 | 0 |
| FXR2_HUMAN | 2/17 | 2 | 0 |
| PPIF_HUMAN | 2/17 | 2 | 0 |
| ERP44_HUMAN | 2/17 | 2 | 0 |
| BZW2_HUMAN | 2/17 | 1 | 0 |
| UBR4_HUMAN | 2/17 | 1 | 0 |
| IF2P_HUMAN | 2/17 | 1 | 0 |
| UBR5_HUMAN | 2/17 | 1 | 0 |
| PREP_HUMAN | 2/17 | 1 | 0 |
| RBM25_HUMAN | 2/17 | 1 | 0 |
| UGGG1_HUMAN | 2/17 | 1 | 0 |
| SNX6_HUMAN | 2/17 | 1 | 0 |
| SPB6_HUMAN | 2/17 | 1 | 0 |
| CSN3_HUMAN | 2/17 | 1 | 0 |

| UniProt ID | Found in palmitoyl-proteomes | Techniques | Times validated |
|---|---|---|---|
| HEM6_HUMAN | 2/17 | 1 | 0 |
| PAK2_HUMAN | 2/17 | 1 | 0 |
| CSN2_HUMAN | 2/17 | 1 | 0 |
| SRC8_HUMAN | 2/17 | 1 | 0 |
| ACADV_HUMAN | 2/17 | 1 | 0 |
| CDC5L_HUMAN | 2/17 | 1 | 0 |
| CBR1_HUMAN | 2/17 | 1 | 0 |
| DDX24_HUMAN | 1/17 | 3 | 0 |
| 2AAB_HUMAN | 1/17 | 3 | 0 |
| BLMH_HUMAN | 1/17 | 3 | 0 |
| CND1_HUMAN | 1/17 | 3 | 0 |
| HDAC2_HUMAN | 1/17 | 3 | 0 |
| IMA4_HUMAN | 1/17 | 3 | 0 |
| LIS1_HUMAN | 1/17 | 3 | 0 |
| PHF6_HUMAN | 1/17 | 3 | 0 |
| SRPK1_HUMAN | 1/17 | 3 | 0 |
| SC24C_HUMAN | 1/17 | 2 | 0 |
| TXNL1_HUMAN | 1/17 | 2 | 0 |
| CYFP2_HUMAN | 1/17 | 2 | 0 |
| TPX2_HUMAN | 1/17 | 2 | 0 |
| PROD_HUMAN | 1/17 | 2 | 0 |
| PP2AA_HUMAN | 1/17 | 2 | 0 |
| TF3C4_HUMAN | 1/17 | 2 | 0 |
| FANCI_HUMAN | 1/17 | 2 | 0 |
| NKRF_HUMAN | 1/17 | 2 | 0 |
| AAAS_HUMAN | 1/17 | 2 | 0 |
| NUP37_HUMAN | 1/17 | 2 | 0 |
| UHRF1_HUMAN | 1/17 | 2 | 0 |
| CCNB1_HUMAN | 1/17 | 2 | 0 |

| UniProt ID | Found in palmitoyl-proteomes | Techniques | Times validated |
|---|---|---|---|
| STAU1_HUMAN | 1/17 | 2 | 0 |
| MOV10_HUMAN | 1/17 | 2 | 0 |
| MP2K1_HUMAN | 1/17 | 2 | 0 |
| CSTF1_HUMAN | 1/17 | 2 | 0 |
| ACL6A_HUMAN | 1/17 | 2 | 0 |
| SMCA1_HUMAN | 1/17 | 2 | 0 |
| KIFC1_HUMAN | 1/17 | 2 | 0 |
| SEPT6_HUMAN | 1/17 | 2 | 0 |
| MAP11_HUMAN | 1/17 | 2 | 0 |
| U3IP2_HUMAN | 1/17 | 2 | 0 |
| NU133_HUMAN | 1/17 | 1 | 0 |
| KIF2C_HUMAN | 1/17 | 1 | 0 |
| KIF5C_HUMAN | 1/17 | 1 | 0 |
| LRBA_HUMAN | 1/17 | 1 | 0 |
| MRE11_HUMAN | 1/17 | 1 | 0 |
| MYG1_HUMAN | 1/17 | 1 | 0 |
| MYO6_HUMAN | 1/17 | 1 | 0 |
| IF2B2_HUMAN | 1/17 | 1 | 0 |
| HECD1_HUMAN | 1/17 | 1 | 0 |
| OGA_HUMAN | 1/17 | 1 | 0 |
| NEST_HUMAN | 1/17 | 1 | 0 |
| GBF1_HUMAN | 1/17 | 1 | 0 |
| OSBP1_HUMAN | 1/17 | 1 | 0 |
| PALLD_HUMAN | 1/17 | 1 | 0 |
| FLII_HUMAN | 1/17 | 1 | 0 |
| FKBP5_HUMAN | 1/17 | 1 | 0 |
| PIPNB_HUMAN | 1/17 | 1 | 0 |
| PNPT1_HUMAN | 1/17 | 1 | 0 |
| PPCE_HUMAN | 1/17 | 1 | 0 |

| UniProt ID | Found in palmitoyl-proteomes | Techniques | Times validated |
|---|---|---|---|
| ECM29_HUMAN | 1/17 | 1 | 0 |
| E41L2_HUMAN | 1/17 | 1 | 0 |
| DYN2_HUMAN | 1/17 | 1 | 0 |
| DPYL5_HUMAN | 1/17 | 1 | 0 |
| DD19A_HUMAN | 1/17 | 1 | 0 |
| RRP44_HUMAN | 1/17 | 1 | 0 |
| SART3_HUMAN | 1/17 | 1 | 0 |
| SCFD1_HUMAN | 1/17 | 1 | 0 |
| SCOT1_HUMAN | 1/17 | 1 | 0 |
| SNR40_HUMAN | 1/17 | 1 | 0 |
| CSTF3_HUMAN | 1/17 | 1 | 0 |
| CSK21_HUMAN | 1/17 | 1 | 0 |
| CPSF1_HUMAN | 1/17 | 1 | 0 |
| SPTN2_HUMAN | 1/17 | 1 | 0 |
| CLAP1_HUMAN | 1/17 | 1 | 0 |
| ZFR_HUMAN | 1/17 | 1 | 0 |
| CDK5_HUMAN | 1/17 | 1 | 0 |
| SUCB2_HUMAN | 1/17 | 1 | 0 |
| SYNE2_HUMAN | 1/17 | 1 | 0 |
| CAN2_HUMAN | 1/17 | 1 | 0 |
| CAN1_HUMAN | 1/17 | 1 | 0 |
| TOM34_HUMAN | 1/17 | 1 | 0 |
| BIRC6_HUMAN | 1/17 | 1 | 0 |
| UBE2O_HUMAN | 1/17 | 1 | 0 |
| UBE3A_HUMAN | 1/17 | 1 | 0 |
| UBP10_HUMAN | 1/17 | 1 | 0 |
| AP3B1_HUMAN | 1/17 | 1 | 0 |
| ANKH1_HUMAN | 1/17 | 1 | 0 |
| UFL1_HUMAN | 1/17 | 1 | 0 |

| UniProt ID | Found in palmitoyl-proteomes | Techniques | Times validated |
|---|---|---|---|
| ACOC_HUMAN | 1/17 | 1 | 0 |
| PFKAM_HUMAN | 1/17 | 1 | 0 |
| PFKAL_HUMAN | 1/17 | 1 | 0 |
| KAP2_HUMAN | 1/17 | 1 | 0 |
| STAT3_HUMAN | 0/17 | 0 | 3 |

**Supplementary table 4:** Out of 264 proteins recovered in WT HAP1 cells labeled with Alk-4 using high confidence settings as described in Figure 4 and Supplementary table, 60 (23%) of proteins were not found in palmitoyl proteomes or validated to be palmitoylated by SWISSPALM. Fifty out of 60 proteins listed here were hits for palmitoylation prediction in SWISSPALM based on CSS-PALM or PalmPred, or had orthologs that were hits in SWISSPalm. In addition, 30 of these proteins were predicted to be palmitoylated by GPS-Lipid(indicated with an *), one protein was predicted to be palmitoylated by CSS-PALM (indicated with an ^), 9 proteins were predicted to be myristoylated by GPS-Lipid (indicated with an +), and 6 proteins were not predicted to be palmitoylated or myristoylated in any database, but contains isoforms and cysteines in TM or cytosolic domain as reported by SwissPalm (indicated by --).

| UniProt ID | Number of isoforms | Maximum number of cysteines | Maximum number of cysteines in TM or cytosolic domain | Predicted to be Spalmitoylated? | Orthologs of this protein have hits in SwissPalm? |
|---|---|---|---|---|---|
| ADRO_HUMAN* | 7 | 10 | 8 | Yes | Yes |
| AFAD_HUMAN* | 6 | 19 | 18 | No | Yes |
| AGM1_HUMAN | 3 | 9 | 8 | Yes | No |
| BLVRB_HUMAN | 1 | 2 | 2 | No | Yes |
| CARF_HUMAN | 1 | 8 | 8 | Yes | No |
| CAZA2_HUMAN | 2 | 4 | 4 | No | Yes |
| CCDC6_HUMAN-- | 1 | 3 | 3 | No | No |

| UniProt ID | Number of isoforms | Maximum number of cysteines | Maximum number of cysteines in TM or cytosolic domain | Predicted to be S-palmitoylated? | Orthologs of this protein have hits in SwissPalm? |
|---|---|---|---|---|---|
| ARMT1_HUMAN-- | 1 | 7 | 7 | No | No |
| CLAP2_HUMAN*+ | 3 | 19 | 14 | Yes | Yes |
| CUL3_HUMAN-- | 3 | 10 | 10 | No | No |
| CUL4A_HUMAN-- | 2 | 9 | 9 | No | No |
| DC1L2_HUMAN* | 2 | 5 | 5 | No | Yes |
| DCTN4_HUMAN | 3 | 16 | 16 | No | Yes |
| TKFC_HUMAN*+ | 2 | 7 | 6 | Yes | Yes |
| DPOD1_HUMAN* | 1 | 26 | 26 | Yes | No |
| DPOLA_HUMAN | 1 | 31 | 31 | Yes | No |
| EFGM_HUMAN* | 2 | 12 | 12 | No | Yes |
| ELP1_HUMAN* | 1 | 29 | 29 | Yes | No |
| EMAL4_HUMAN*+ | 2 | 19 | 19 | No | No |
| EFL1_HUMAN* | 2 | 23 | 23 | Yes | No |
| FTO_HUMAN+ | 4 | 14 | 14 | Yes | No |
| GAPD1_HUMAN* | 6 | 26 | 26 | Yes | No |
| GRDN_HUMAN | 5 | 12 | 12 | No | Yes |
| GYS1_HUMAN | 2 | 14 | 14 | No | Yes |
| HCFC1_HUMAN* | 4 | 35 | 35 | Yes | Yes |
| IDH3B_HUMAN | 3 | 7 | 5 | Yes | Yes |
| IF4G3_HUMAN* | 4 | 20 | 20 | Yes | Yes |
| INT1_HUMAN* | 1 | 39 | 39 | Yes | No |
| IQGA2_HUMAN* | 3 | 10 | 10 | Yes | Yes |
| JIP4_HUMAN+ | 6 | 15 | 14 | Yes | No |
| KDM1A_HUMAN-- | 2 | 9 | 9 | No | No |
| KIF2A_HUMAN | 5 | 12 | 12 | Yes | No |
| KIF4A_HUMAN* | 2 | 28 | 28 | Yes | Yes |

| UniProt ID | Number of isoforms | Maximum number of cysteines | Maximum number of cysteines in TM or cytosolic domain | Predicted to be S-palmitoylated? | Orthologs of this protein have hits in SwissPalm? |
|---|---|---|---|---|---|
| KLC2_HUMAN | 2 | 7 | 7 | No | Yes |
| KPRA_HUMAN | 2 | 5 | 5 | No | Yes |
| KS6A1_HUMAN+ | 4 | 8 | 7 | Yes | Yes |
| MCMBP_HUMAN* | 3 | 15 | 15 | Yes | Yes |
| MVD1_HUMAN* | 1 | 9 | 9 | No | No |
| NEK9_HUMAN* | 1 | 31 | 31 | Yes | No |
| NIPS1_HUMAN* | 1 | 3 | 3 | Yes | Yes |
| NRDC_HUMAN* | 2 | 18 | 18 | Yes | No |
| NUP85_HUMAN+ | 3 | 15 | 15 | Yes | Yes |
| PLCB3_HUMAN* | 2 | 14 | 14 | No | Yes |
| PLCG1_HUMAN*+ | 2 | 23 | 23 | Yes | Yes |
| PLSI_HUMAN | 1 | 7 | 7 | Yes | Yes |
| PTN11_HUMAN | 3 | 10 | 10 | No | Yes |
| RAD50_HUMAN* | 3 | 18 | 18 | Yes | No |
| RB_HUMAN^ | 1 | 15 | 15 | No | No |
| SAHH2_HUMAN* | 2 | 19 | 19 | Yes | Yes |
| SAMH1_HUMAN* | 4 | 13 | 13 | No | Yes |
| SC23B_HUMAN | 1 | 17 | 17 | Yes | Yes |
| SMHD1_HUMAN+ | 3 | 30 | 30 | Yes | No |
| SNW1_HUMAN-- | 1 | 1 | 1 | No | No |
| SUGP2_HUMAN | 3 | 17 | 16 | Yes | No |
| FAKD4_HUMAN | 2 | 8 | 8 | Yes | Yes |
| TLN2_HUMAN* | 1 | 41 | 41 | Yes | Yes |
| TXLNG_HUMAN* | 2 | 9 | 9 | No | No |
| VP13C_HUMAN | 4 | 41 | 41 | Yes | Yes |
| WNK1_HUMAN* | 6 | 26 | 16 | Yes | Yes |

| UniProt ID | Number of isoforms | Maximum number of cysteines | Maximum number of cysteines in TM or cytosolic domain | Predicted to be S-palmitoylated? | Orthologs of this protein have hits in SwissPalm? |
|---|---|---|---|---|---|
| XPOT_HUMAN* | 1 | 22 | 22 | No | Yes |
